# Differential Gene Expression Analysis of a Small Colony Variant of *Escherichia coli* K-12

**DOI:** 10.1101/413468

**Authors:** Anthony W. Paratore, Victors Santos, Stergios Bibis, Irvin N. Hirshfield

## Abstract

As persistent residents of planktonic bacterial cultures, small colony variants (SCVs) constitute a slow-growing subpopulation with atypical colony morphology and unusual biochemical characteristics that, in the case of clinical isolates, cause latent and recurrent infections. We propose a novel blueprint for the formation of *E. coli* SCVs through DNA microarray analysis, coupled with complete genome sequencing and verification by qRT-PCR. While others have used DNA microarrays to study quorum sensing in *E. coli* SCVs, our work represents the first proposal for a combination of novel mutations, amplified by a differential shift in expression of select gene groups that work in concert to establish and maintain the SCV phenotype. This combination of genetic and expression events fall under selective pressure, leading to unequal fitness in our strain, SCV IH9 versus its parental strain, BW7261 (a MG1655 descendant). We hypothesize that this combination of events would ordinarily be lethal for bacteria, but instead confers a survival advantage to SCV IH9 due to its slow growth and resistance to acidic and oxidative stress challenges.

## Introduction

Resistance to environmental stresses is the paradigm of evolutionary fitness in the microbial world. To counter environmental stresses, bacteria have developed resistance to the oxidative damage they encounter in macrophages, antibiotics encountered in mammalian hosts, and pH or acid shifts [1-3]. Natural selection has produced survival mechanisms in bacteria such as persister cells, biofilms and small colony variants (SCVs) [4-6]. As true phenotypic variants and persistent residents of planktonic bacterial cultures, SCVs constitute a slow-growing subpopulation with atypical colony morphology and unusual biochemical characteristics that, in the case of clinical isolates, cause indolent and recurrent infections [7-10].

SCVs were first described in 1910 by Jacobsen, who found abnormally small colonies in a population of wild type *Salmonella enterica* [11]. In subsequent years, SCVs of *S. aureus, S. epidermidis, P. aeruginosa, V. cholerae, S. marcescens, N. gonorrhoeae* and others have been isolated and identified [12]. SCVs of the above mentioned bacteria all display similar morphological colony characteristics such as small size, smooth (glossy) convex surface morphology, and slow growth [13].

Our knowledge and understanding of physiological and molecular mechanisms that define SCVs has come predominantly from the work of Richard Proctor the past three decades. Proctor and colleagues report SCVs of *S. aureus* as persistent subpopulations within normal planktonic communities that are ordinarily out-competed by the wild type members of the culture [14]. Small colony variants of *S. aureus* (described as auxotrophs) exhibit a genetic mutation rendering them unable to synthesize one of three important metabolites: hemin, thymidine or menadione. SCVs defective in hemin and menadione synthesis are classified as electron transport defective SCVs due to their hindered production of ATP as a result of said mutation [14]. This lack of adequate energy production accounts for their reduced colony size and unique biochemical properties such as acid resistance.

SCVs have been isolated in patients with underlying chronic/persistent infections in skin and soft tissue. After clearing their infection with the wild-type strain, SCVs have been found to persist in patients for months or years [15]. Clinicians frequently report bloodstream infections due to *S. aureus* SCVs following endocardial pacemaker implantation [16]. The persistence of SCVs in living tissue presents an abnormal survival advantage compared to the normal wild type phenotype despite the presence of immune cells.

Despite much work focused on understanding the physiological properties of SCVs, very little is known concerning the molecular mechanisms leading to SCV formation in non-clinical *E. coli* SCVs. Much of the work on *E. coli* SCVs has focused on understanding its physiology but our understanding of molecular mechanisms involved in SCV formation remains limited. *E. coli* SCVs are typically identified by their morphology but a molecular profile of an *E. coli* SCV is incomplete at best.

In this study, the SCV analyzed (SCV IH9) displays an altered phenotype consisting of slow growth, propensity to form biofilms, markedly enhanced survival in low pH and upon exposure to agents of oxidative damage [17]. Microarray studies were performed to better understand the basis of the SCV phenotype and to identify genes and/or gene pathways that could contribute to this altered phenotype. In addition, qRT-PCR was used to verify the DNA microarray and DNA sequencing of the entire genome was conducted to explore single nucleotide polymorphism (SNP) variations.

In this report, it is shown that a SCV of *E. coli* will display differential gene expression as a strategy to express traits that will allow it to endure survival challenges. In addition, our discussion will focus on the evolutionary importance of genome-wide differential gene expression of an *E. coli* SCV.

## Materials and Methods

### Bacterial strains

The *E. coli* K-12 strains of Escherichia coli used in this study are listed in Table 1.

**Table 1.** *E. coli* K-12 strains (descendants of MG1655)

| Strain | Genotype |
| --- | --- |
| BW7261 | Parental strain of IH9* |
| SCV IH9 | Small colony variant of BW7261<sup>§</sup> |

\* *E. coli* Genetic Stock Center, Yale University
<sup>§</sup> Hirshfield Laboratory, St. John's University. SCVs isolated by acid resistance.

### Media and cell culture techniques

Cells were grown aerobically in Luria-Bertani broth. Five ml starter cultures were grown to log phase in test tubes by inoculating with cells of a fresh bacterial colony grown on LB agar. For all experiments, 200 ml of bacterial culture were incubated in 1000 ml growth flasks in a controlled environment incubator shaker (200 rpm, 37°C) until cells reached log phase. Growth of cultures was periodically monitored by measuring the optical density (OD) at 580 nm using a spectrophotometer (Carolina Digital Spectrophotometer Model # 653303). Cells were grown to log phase (approximately 1 x 10^8^ cells/ml). Cells were harvested by centrifugation in a Beckman Avanti™ J-25 centrifuge for 20 minutes at 8,000 RPM (7728 RCF).

### RNA isolation and extraction

SCV IH9 and BW7261 were grown to log phase and total RNA was extracted using an Ambion RiboPure™ Bacteria Kit (Applied Bioscience). Genomic DNA was eliminated by RNase-free DNase I during the isolation. Each RNA sample was quantified spectrophotometrically for quality and quantity. The RNA was then eluted into elution buffer and stored at −80°C. RNA ranged in concentration from 300 - 900 ng/µl. The 260/280 ratio for all samples ranged from 1.8 – 2.0.

### DNA Microarray

Total RNA samples were shipped to a core facility (MoGene, LLC – St. Louis, MO) for analysis. MoGene purified the samples of solvent contamination from the RNA extraction kits to their optimal purity. cDNA synthesis and labeling of cDNA via reverse transcription was performed at MoGene (SCV IH9 cDNA was labeled with Cy5, a green fluorescent dye exciting at 650 nm and BW7261 cDNA was labeled with Cy3, a red fluorescent dye exciting at 550 nm).

### DNA extraction for sequencing

For sequencing purposes, genomic DNA was extracted from the strains listed in Table 1 using a Sigma-Aldrich ® Bacterial Genomic Miniprep Kit. DNA was eluted with buffer and quantified spectrophotometrically for quality and quantity. All samples had concentrations greater than 200 ng/µl and 260/280 ratios ranging from 1.73 - 2.16.

### DNA sequencing

Illumina® Inc. sequencing was performed by The University at Buffalo Next-Generation Sequencing and Expression Analysis Core Facility (UB Next-Gen Core). Sequencing services were performed using the Roche/454 Genome Sequencer FLX and Illumina HiSeq 2000 platforms. All strains listed in Table 1 were sequenced. Illumina generated the DNA sequencing libraries for each strain and prepared all samples for sequencing. Genomes were sequenced with a 50-cycle, single read sequencing experimental design that provided in excess of 160 million reads per flow cell lane. As *E. coli*’s genome is less than 5 megabases in length, the sequencing coverage was greater than 800X, allowing greater confidence in SNPs found. Geneious® Bioformatics Software version 10.1 (created by Biomatters LTD) was used for analysis, interpretation, and application of molecular sequence data.

### Quantitative Reverse Transcription Polymerase Chain Reaction

qRT-PCR, which uses RNA as a starting nucleic acid, begins with the reverse transcription of RNA into complementary DNA (cDNA) by reverse transcriptase from total RNA or messenger RNA (mRNA) with the cDNA then used as the template for the PCR reaction. This experiment was performed by RUCDR-Infinite Biologics (Piscataway, NJ) based out of Rutgers University, who handled all steps excluding the growth of BW7261 and SCV IH9. Briefly, RUCDR performed RNA extraction and processing, assay design and qRT-PCR, etc.

## Results

Coupling the results of the DNA microarray and genome sequencing we propose a unique series of genetic and phenotypic events that establish and maintain the SCV phenotype in *E. coli* communities.

### Gene expression in SCV IH9 is markedly different from wild type gene expression

DNA microarray studies revealed several critical gene groups that are over-expressed or repressed in SCV IH9. Key to our study is the pattern of expression related to genes involved in iron transport, colanic acid production, anaerobic-aerobic regulation, lipopolysaccharide formation and LPS composition and general stress-response genes. These groups of genes were chosen due to their role in contributing to survival in other organisms (Fig. 1).

**Fig. 1.**
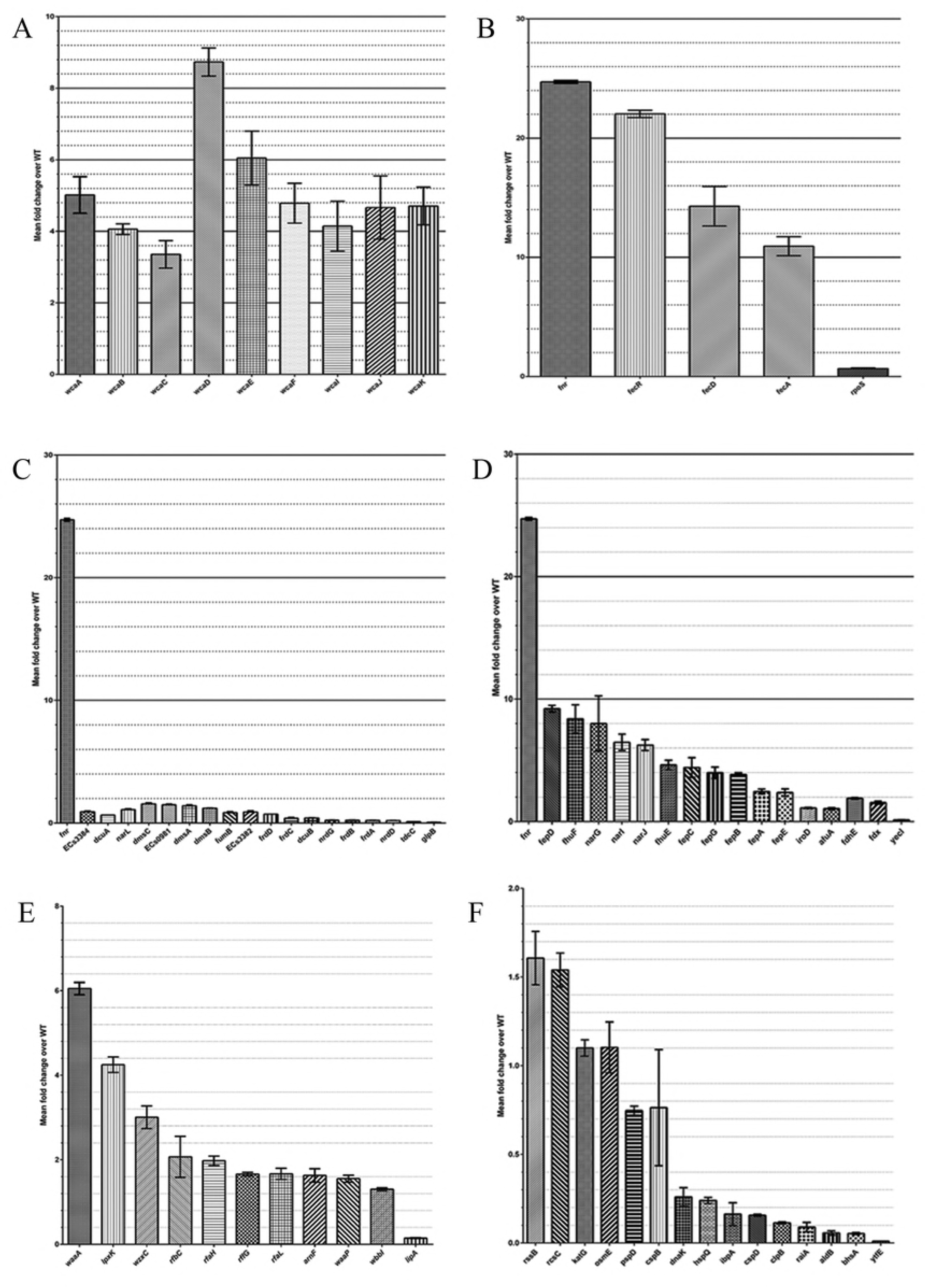
SCV IH9 vs. BW7261 Microarray data. **A**| *wca* genes, involved in the production of colanic acid, are differentially over-expressed in IH9. **B**| Expression of *fnr*, a dual transcriptional regulator and global transcription factor for anaerobic growth is shown alongside several over- expressed *fec* genes (*fecR, fecD, fecA*). **C**| Anaerobic genes differentially expressed across the genome. **D**| Ferric genes. **E**| Genes involved in lipopolysaccharide formation and LPS composition. **F**| Stress-response genes. Genes with a mean fold change ≈ 1 display normal expression, mean fold change < 1 indicates gene repression, mean fold change > 1 indicates over expression of gene. DNA microarray replicated data were combined, and the mean-fold change are presented. All data were calculated from experimental results obtained from at least three independent cultures.

We propose this pattern of differential gene expression is a novel requirement of select *E. coli* SCVs. Differential expression of unique gene groups (*e.g.* ferric genes) demonstrates SCV IH9’s stress response despite the fact it was grown in regular media under aerobic conditions.

### Nonsense mutations may contribute to the SCV phenotype

Currently, one published article highlights complete genome sequencing of *E. coli* SCVs [18]. Our study identifies SNPs in a very small percentage of *E. coli*’s genes that may be critical for the establishment and maintenance of the SCV phenotype. We have identified four SNP-containing genes in SCV IH9 that possess mutations (SNPs) that carry severe consequences for the gene and protein *(cadB, glcD, MmuP*, and *ompF*).

### *cadB* gene

The CadB protein is part of the lysine-dependent acid resistance system which confers resistance to weak acids produced during carbohydrate fermentation under conditions of anaerobiosis [19] and may be regulated by extracellular pH [20]. The *cadB* gene (Fig. 2) in both BW7261 and SCV IH9 displays a substitution at nucleotide 718 (C → T). This is not unexpected as SCV IH9 was isolated from BW7261 and shows only slight genetic divergence from it. This substitution results in a premature stop codon in the CadB protein at codon 240 (glutamine → STOP). The protein is normally 444 amino acid residues long; the CadB protein in both BW7261 and SCV IH9 is 240 amino acids in length. BW7261 and SCV IH9 both possess a SNP that severely truncates the normal CadB protein (the truncated protein is 240 amino acids in length but wild type CadB protein is 444 amino acids long) (Fig. 3). The ability of the *cadB* gene to produce functional protein is compromised reducing CadB’s ability to import lysine. For *E. coli*, lysine provides protection during anaerobic starvation and one study showed that a *cadBA* deletion completely reverses this effect [21].

**Fig. 2.**
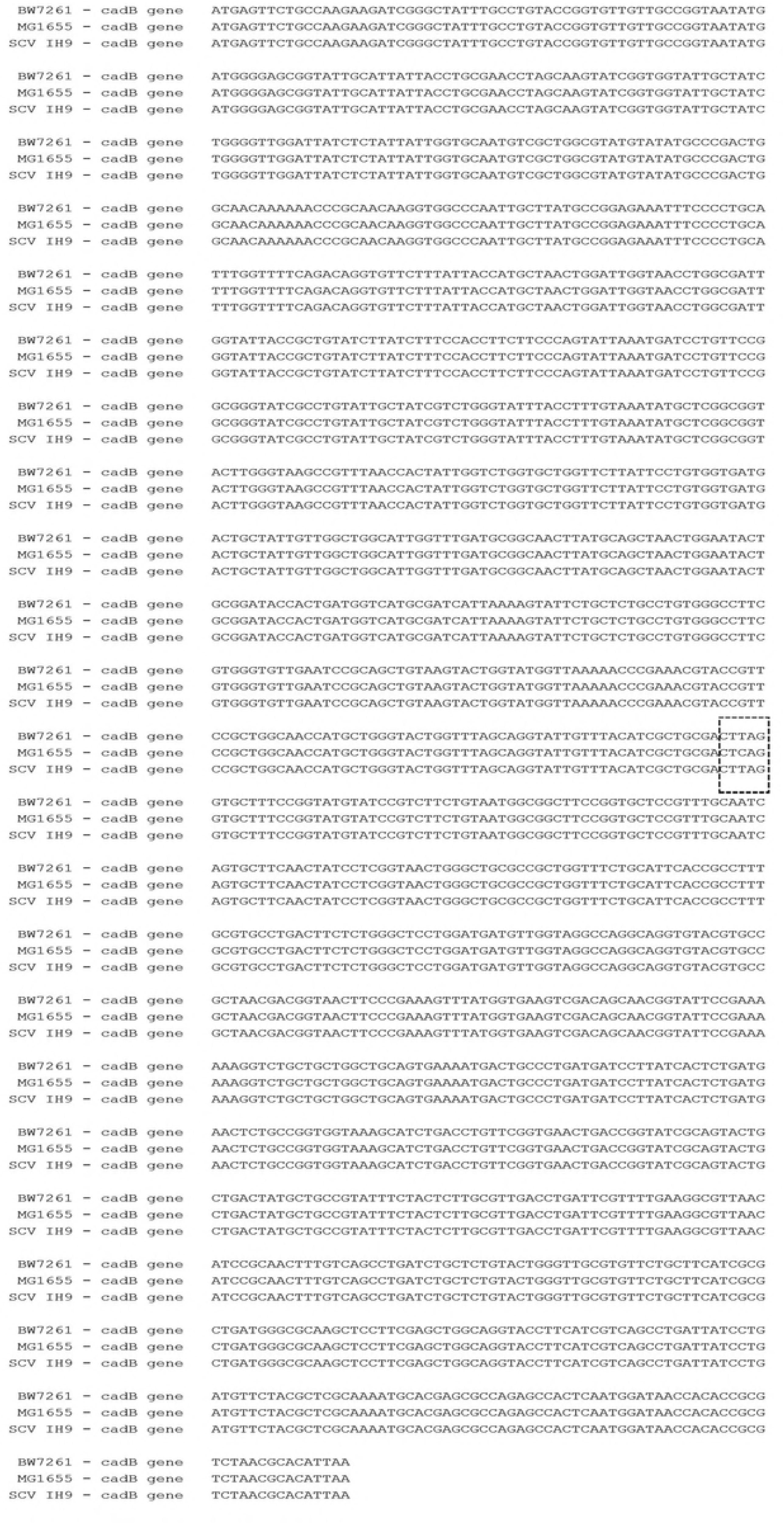
*cadB* gene nucleotide alignment of SCV IH9 vs. BW7261. (with respect to reference genome MG1655). SNP is highlighted in dotted box.

**Fig. 3.**
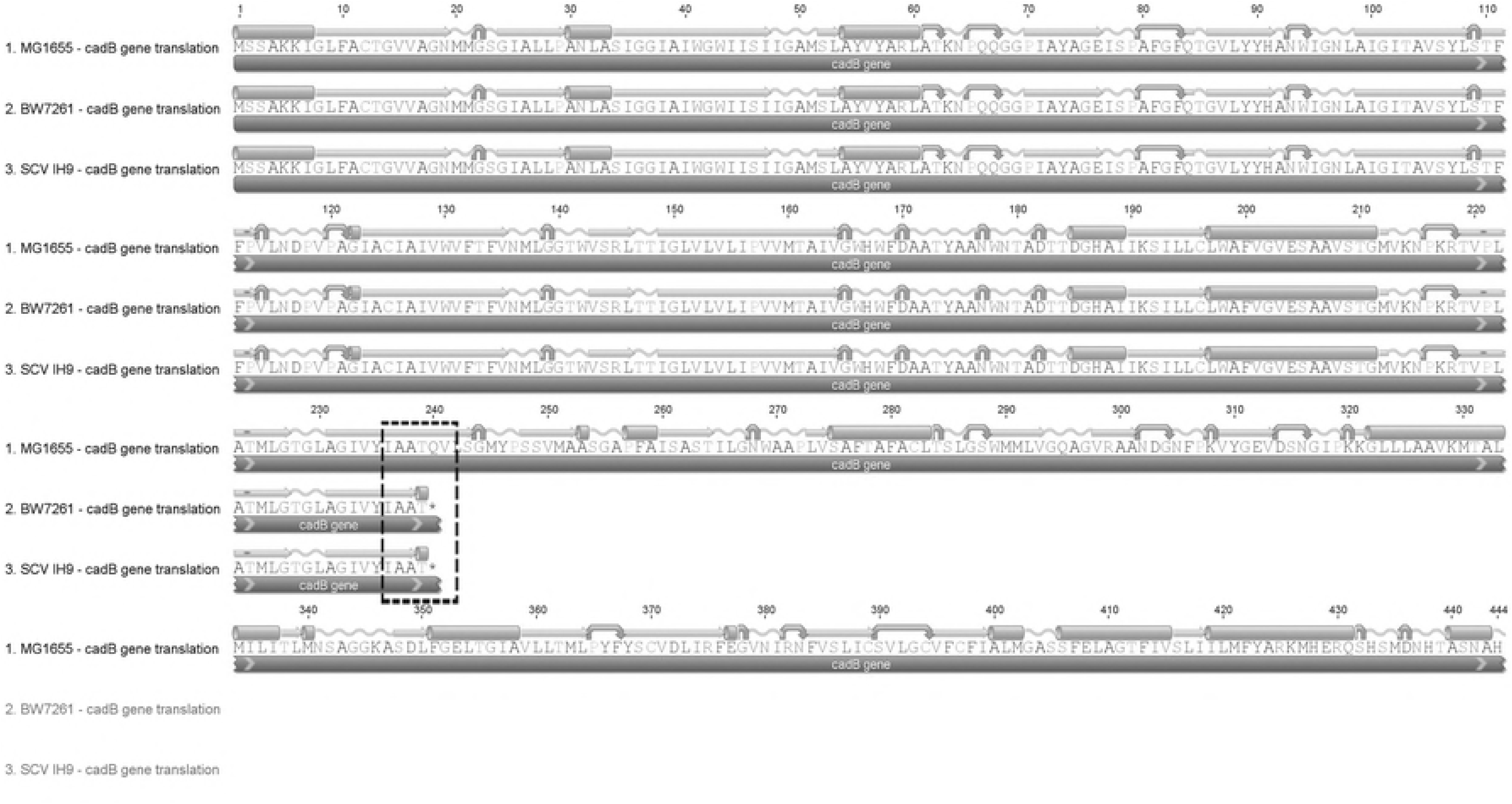
Protein alignment for CadB in SCV IH9 vs. BW7261. (with respect to reference genome MG1655). Premature stop codon is highlighted in dotted box.

### *glcD* gene

*GlcD* encodes a component of the glycolate oxidase complex (GlcD protein), which catalyzes the first step in the utilization of glycolate as a source of carbon [22]. In SCV IH9, the SNP at nucleotide 587 (C → A) results in a premature stop codon in the GlcD protein at amino acid codon 196 (Fig. 4). In SCV IH9, this results in a GlcD protein of 195 amino acids in length whereas wild type GlcD is normally 499 amino acid residues in length (Fig. 5). Older studies have demonstrated that glycolate oxidase is composed of several components (GlcD being one component) and any insertional mutations that silence either *glcD, glcE*, or *glcF* would abolish the enzyme’s activity [23].

**Fig. 4.**
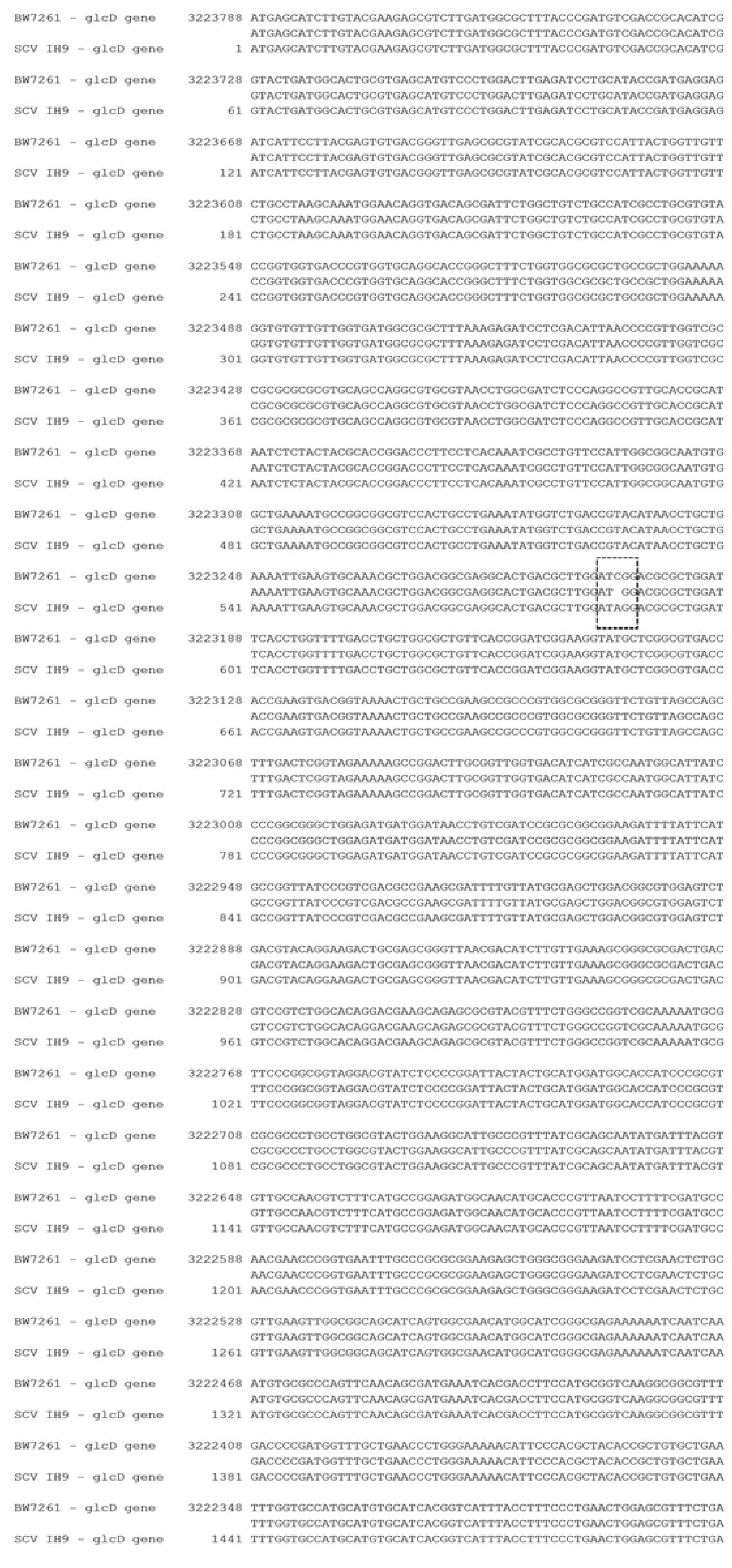
*glcD* gene nucleotide alignment of SCV IH9 vs. BW7261. (with respect to reference genome MG1655).

**Fig. 5.**
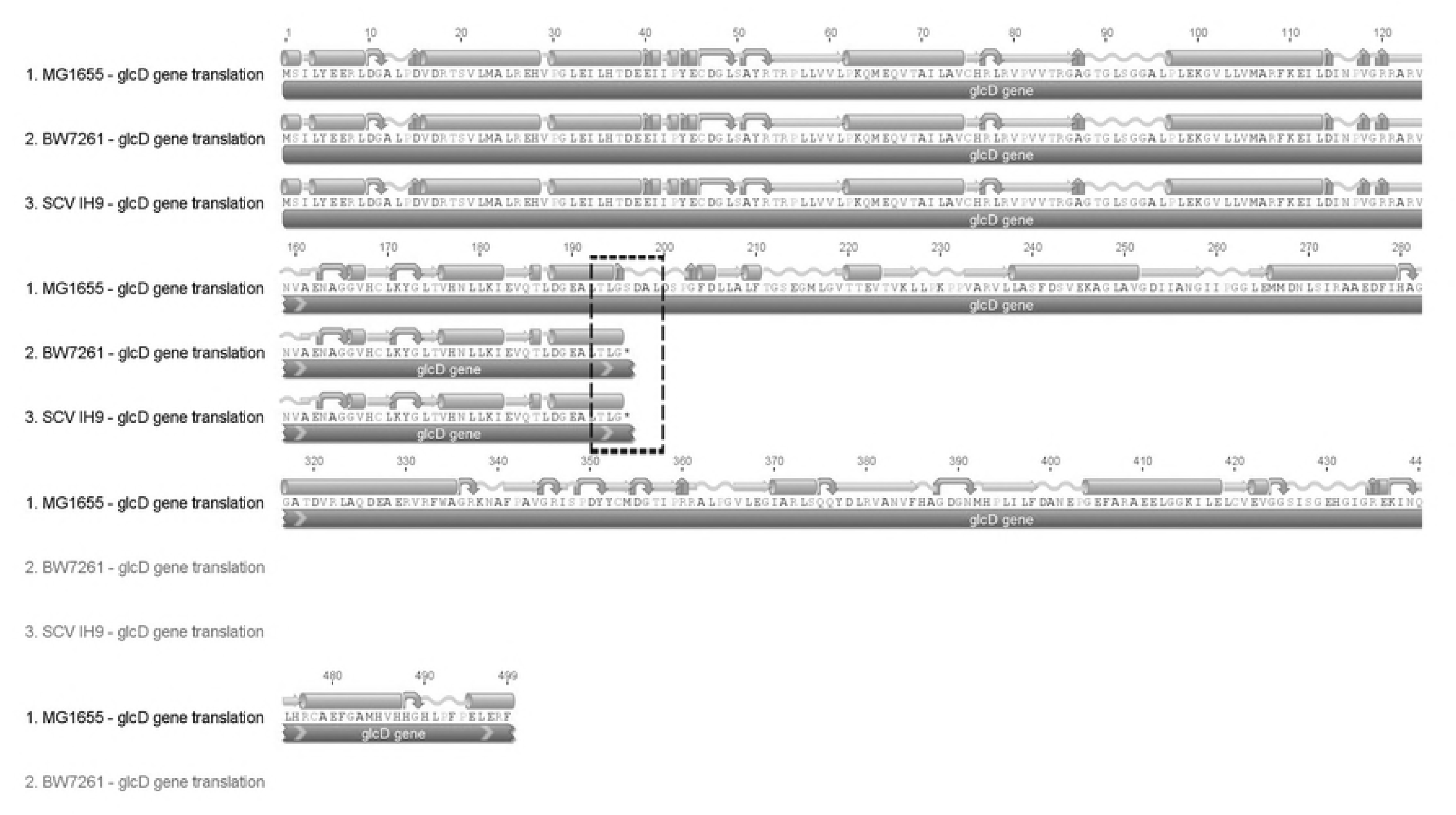
Protein alignment for GlcD in SCV IH9 vs. BW7261 showing residues near codon 196. (premature stop codon).

***MmuP*** gene

*MmuP* (“**m**ethyl**m**ethionine **u**tilization”) encodes a putative S-methylmethionine transporter and mutants with in-frame deletions lack the ability to utilize S-methylmethionine as a source of methionine [24]. The *mmuP* gene (Fig. 6) bears a deletional mutation at nucleotide 185-186 (TT → C). In SCV IH9, the MmuP protein has an amino acid substitution at codon 62 (valine → alanine) as a result of the deletion mutation (Fig. 7). Consequently, the amino acid residues 63 to 100 are all mutated, because of the frame shift generated by the deletional mutation. Moreover, a premature stop codon is introduced at amino acid residue 101, truncating the remaining 367 amino acids of the protein. Any activity of MmuP (*e.g*., methionine storage, methionine biosynthesis) is believed to be abolished in SCV IH9.

**Fig. 6.**
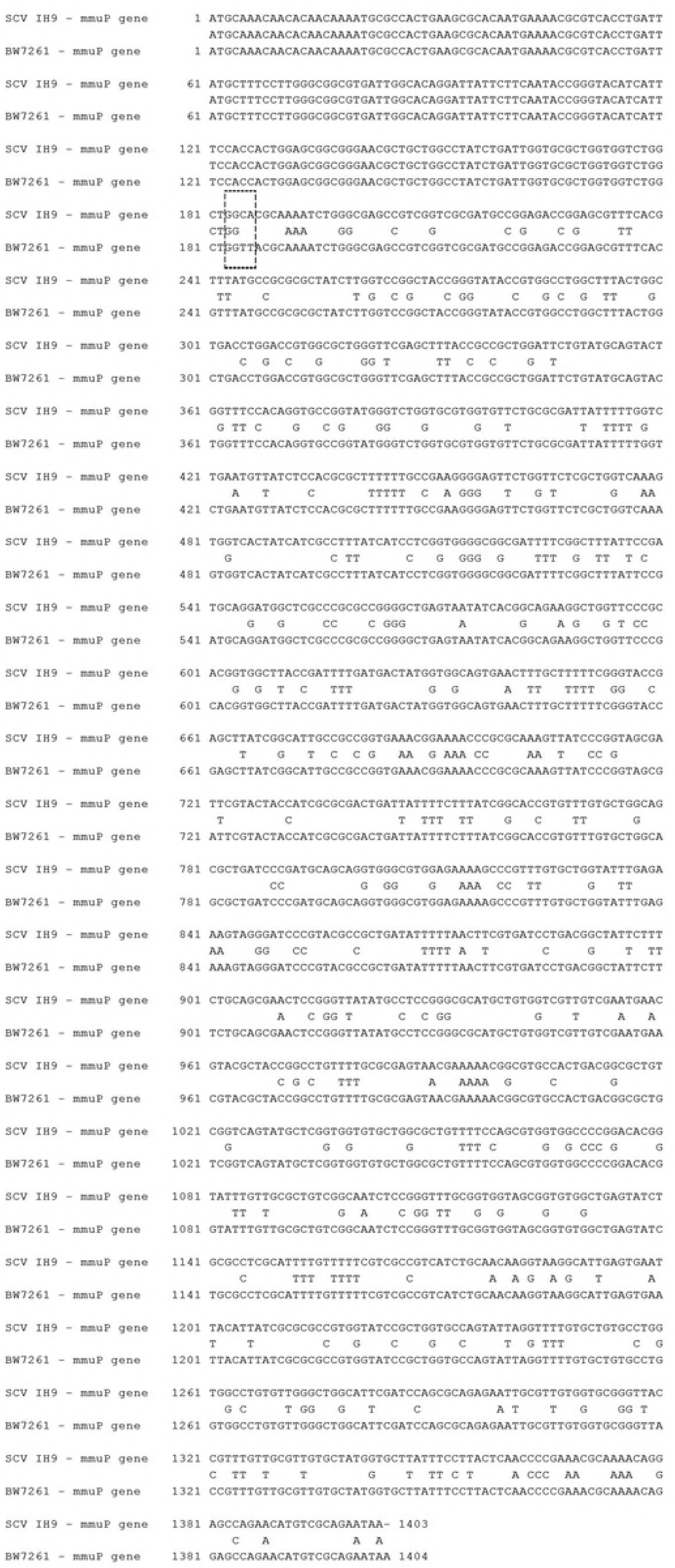
*MmuP* gene nucleotide alignment of SCV IH9 vs. BW7261. Nonsense SNP is highlighted in dotted box.

**Fig. 7.**
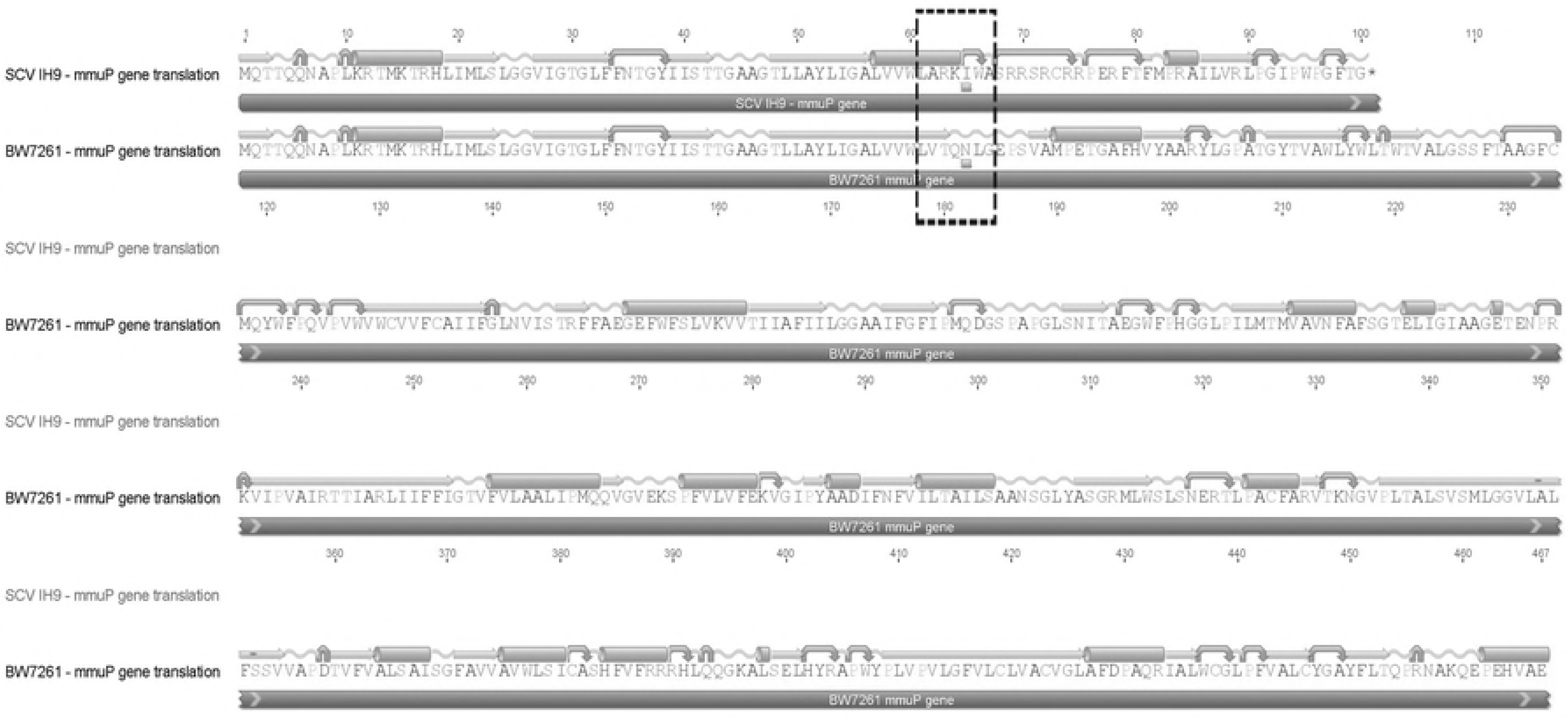
Protein alignment for MmuP in SCV IH9 vs. BW7261. Premature stop codon is highlighted in dotted box.

### *ompF* gene

Finally, the *ompF* gene encodes an outer membrane porin that permits solutes such as sugars, ions, and amino acids to enter or exit the cell [25]. The *ompF* gene of SCV IH9 (Fig. 8) contains a substitution at nucleotide 673 (C → T) which results in a premature stop codon in the OmpF protein at amino acid codon 225 truncating the protein’s remaining 138 amino acids (Fig. 9). OmpF is believed to be the main pathway for β-lactam antibiotics to permeate the cell; for SCV IH9 this may translate into antibiotic resistance.

**Fig. 8.**
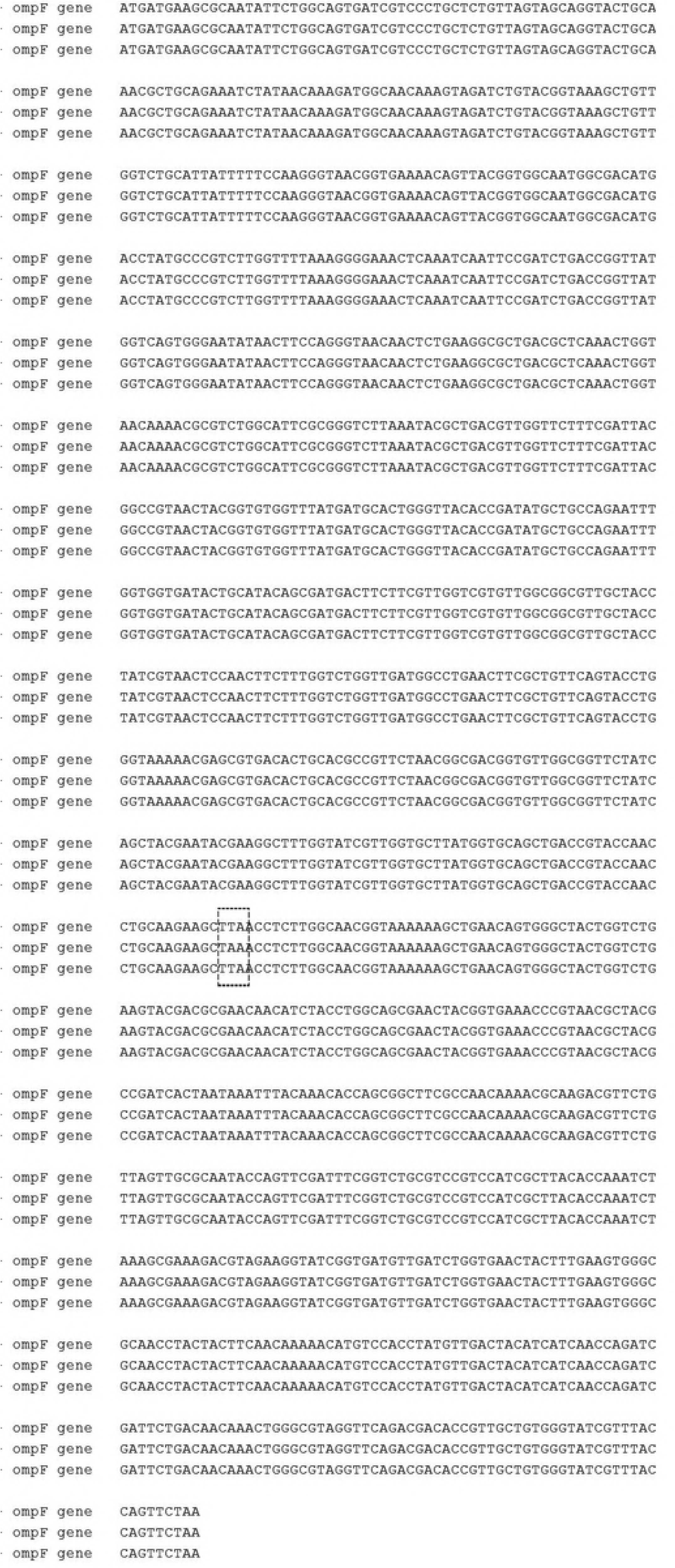
*OmpF* gene nucleotide alignment of SCV IH9 vs. BW7261. (with respect to reference genome MG1655). SNP is highlighted in dotted box.

**Fig. 9.**
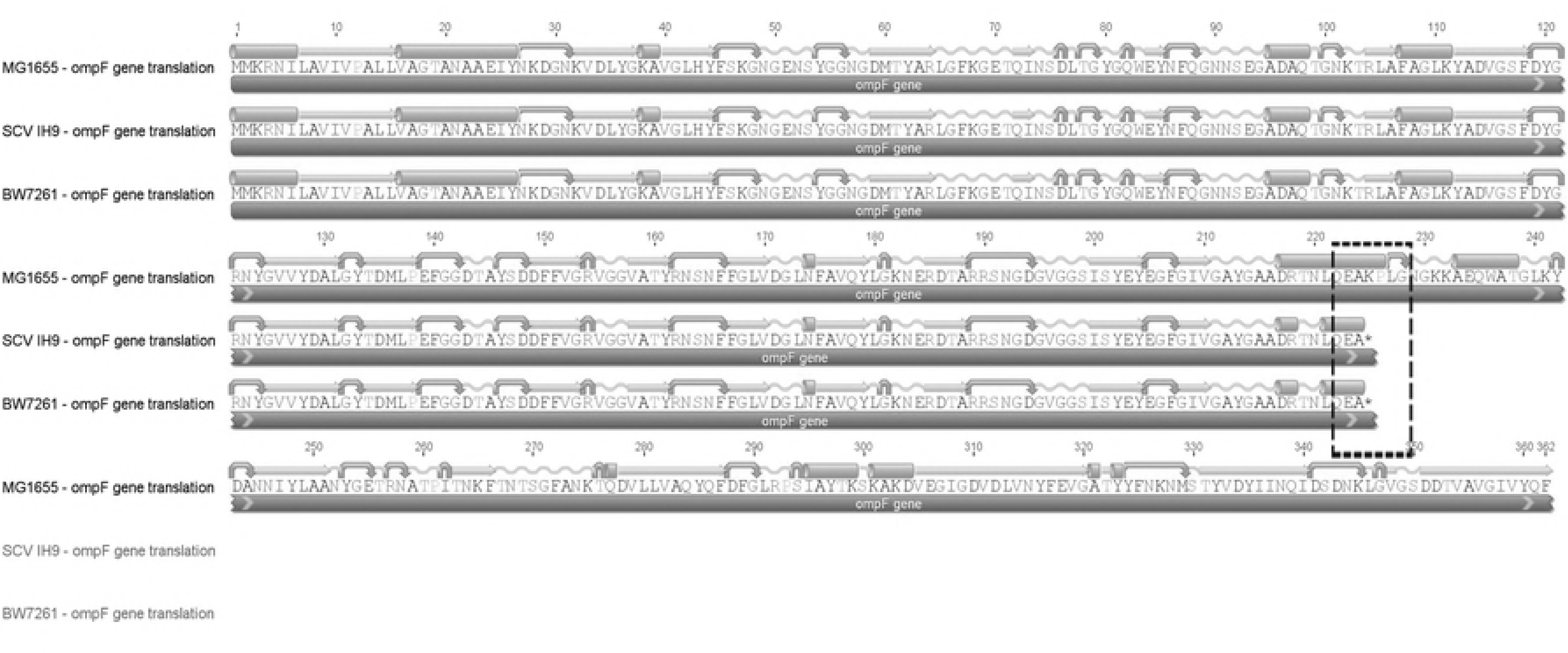
Protein alignment for OmpF in SCV IH9 vs. BW7261. Premature stop codon is highlighted in dotted box.

While these four mutated genes (and proteins) may not engender a classic stress response state in SCV IH9 (*e.g*., cold shock or heat shock), it may leave SCV IH9 incapable of metabolizing glycolate, methionine, and mounting a select stress response (*e.g.,* colanic acid for biofilm formation).

### SCV IH9 SNPs may influence gene expression of important genes

There are several SNPs (*grxB, lpxK, torD,* etc.) that are found only in SCV IH9 but not BW7261 (Table 2). None of these SNPs results in truncations of the protein product of their respective genes. The missense substitutions may be tolerable (resulting in no significant change to amino acid sequence and no loss-of-function for the protein) or intolerable (where even a single change in amino acid renders a protein mutant or nonfunctional). Our study analyzes protein structure utilizing Geneious^®^’ protein domain prediction software plug-in.

**Table 2.**
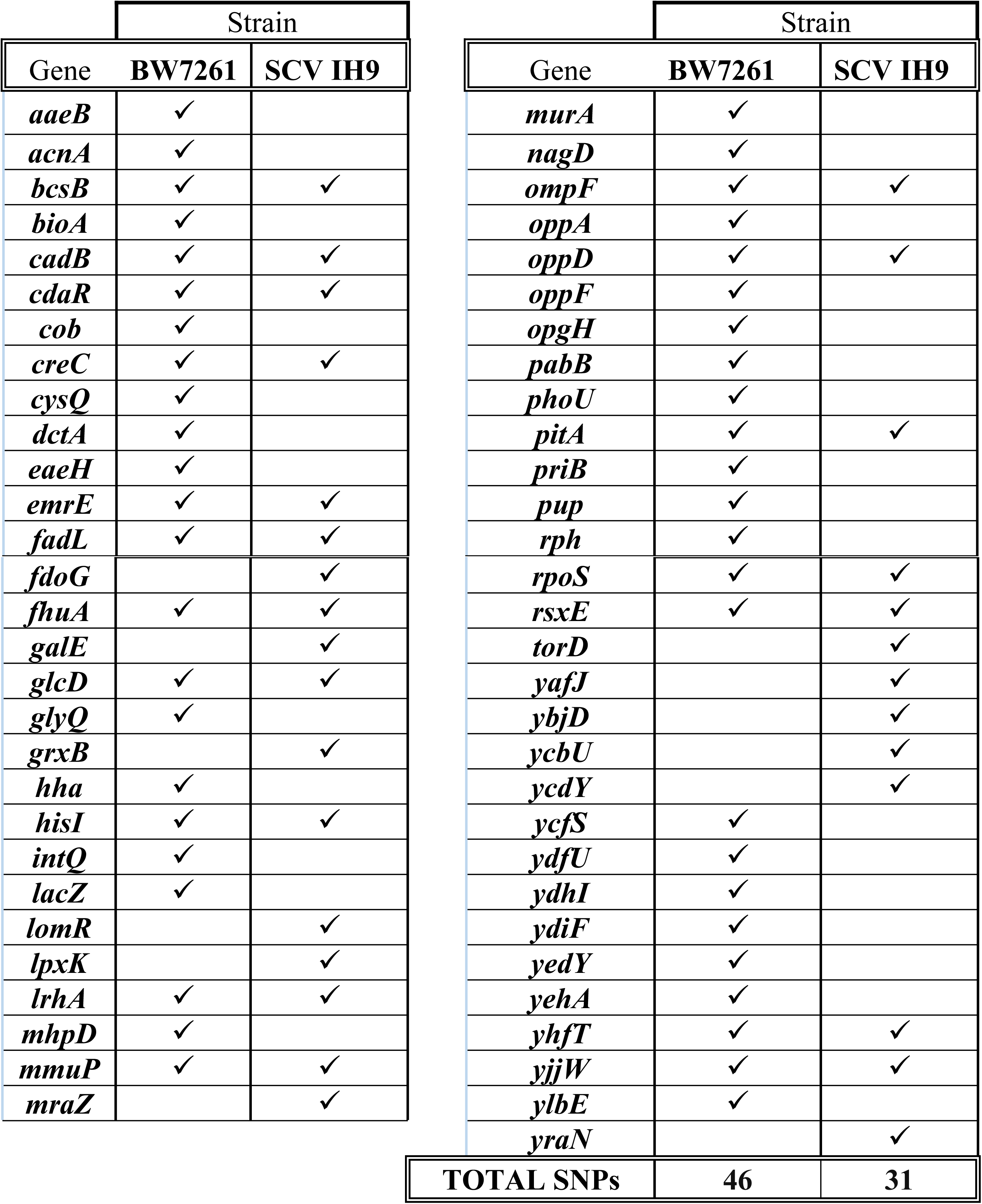
Comparison of SNPS (SCV IH9 vs BW7261).

*GrxB* encodes three different glutaredoxins that catalyze the reduction of disulfides via reduced glutathione. *E. coli* has three glutaredoxins (Grx1, Grx2, and Grx3) which function as cofactors permitting intracellular redox reactions [26].

Grx2 is *E. coli*’s most abundant glutaredoxin that reduces cytosolic protein disulfides and stimulates the reconstitution of the [4^Fe^-4^S^] cluster of FNR [27]. SCV IH9 possesses a SNP in the *grxB* gene at nucleotide 499 (A → G) (Fig. 10) that causes an amino acid substitution at residue 167 (lysine → glutamic acid) (Fig. 11). While the consequences of this change are presently unknown, the amino acid substitution is significant because lysine is a basic and positively charged amino acid while glutamic acid is acidic and negatively charged. The sequencing software used in this study (Geneious®) predicts that this substitution results in a shorter coil and longer alpha helix immediately downstream of this site. This gene (and protein) should be further explored because of the relationship that exists between GrxB and Fnr.

**Fig. 10.**
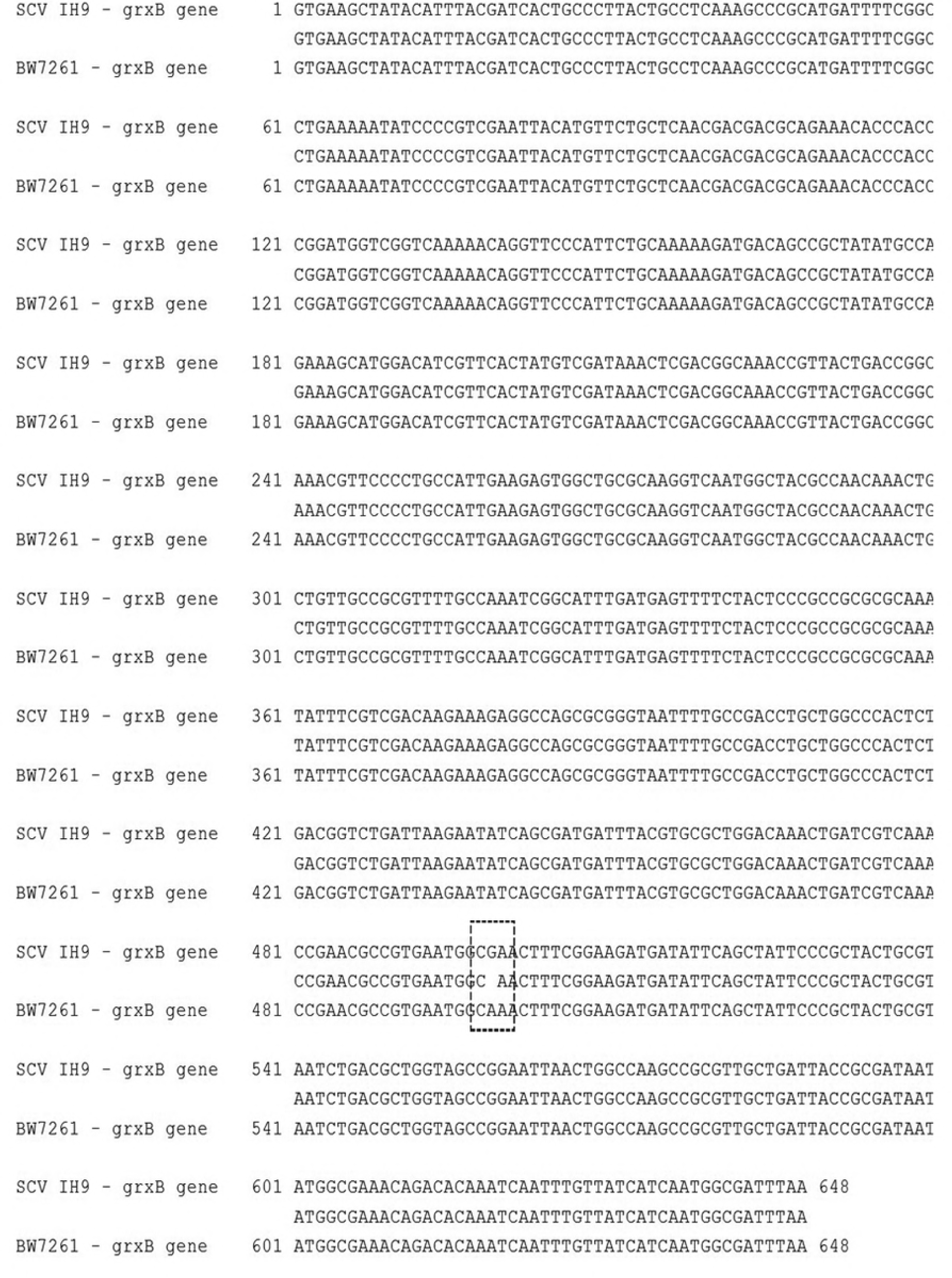
*GrxB* gene nucleotide alignment of SCV IH9 vs. BW7261. SNP is highlighted in dotted box.

**Fig. 11.**
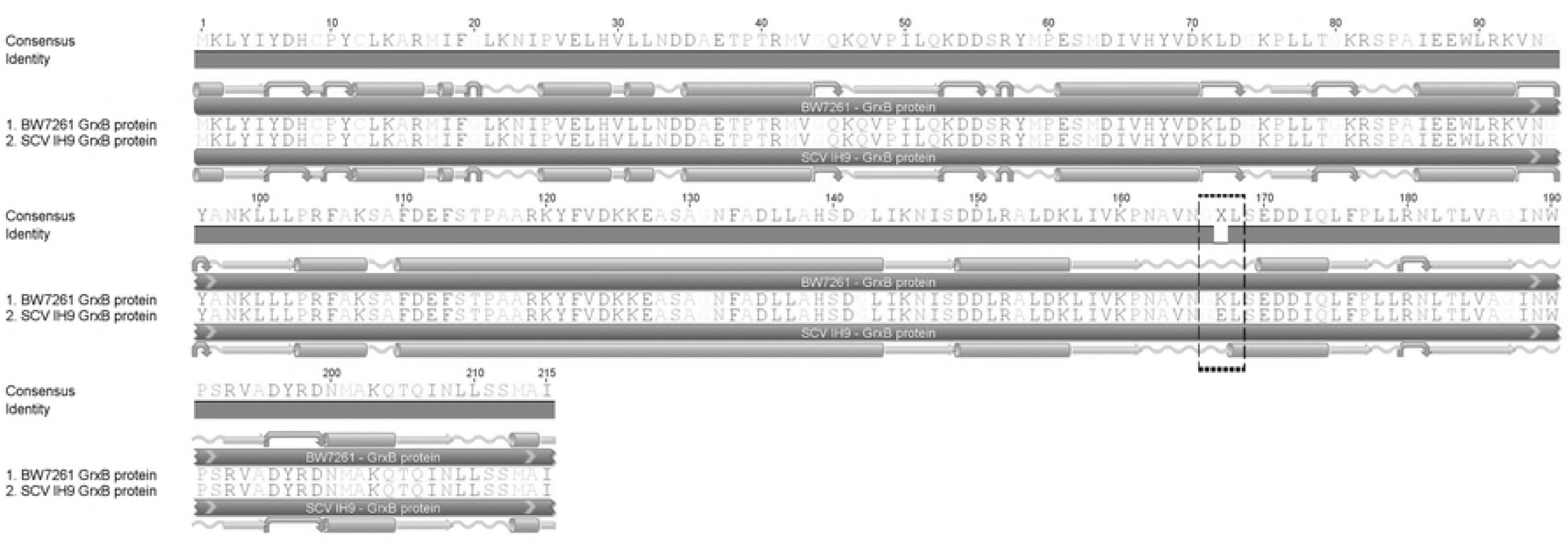
Protein alignment for GrxB in SCV IH9 vs. BW7261. Amino acid substitution is highlighted in dotted box.

*LpxK* encodes a lauroyl acyltransferase that catalyzes the sixth step in lipid A biosynthesis, forming the most immediate lipid A precursor [28]. *LpxK*, was first identified as *orfE* in *E. coli* and has been demonstrated to be critical to cell survival as mutants lacking this gene are not viable [29]. *lpxK* is overexpressed in SCV IH9 (mean fold change of 4.18). SCV IH9 possesses a SNP in the *lpxK* gene at nucleotide 472 (A → G) (Fig. 12) that causes an amino acid substitution at residue 158 (aspartic acid → asparagine). While the consequence of this change are presently unknown, the amino acid substitution is significant because aspartic acid is acidic and negatively charged while asparagine is a neutral amino acid. Sequencing software predicts that this substitution changes the coils at this residue (and adjacent amino acids 155 –157) into a four-residue alpha helix at residues 155 - 158 (Fig. 13). To date, there is one study highlighting this precise mutation (D → N at residue 158) in *E. coli* (strain DH10B) but no phenotypic changes were detected [30].

**Fig. 12.**
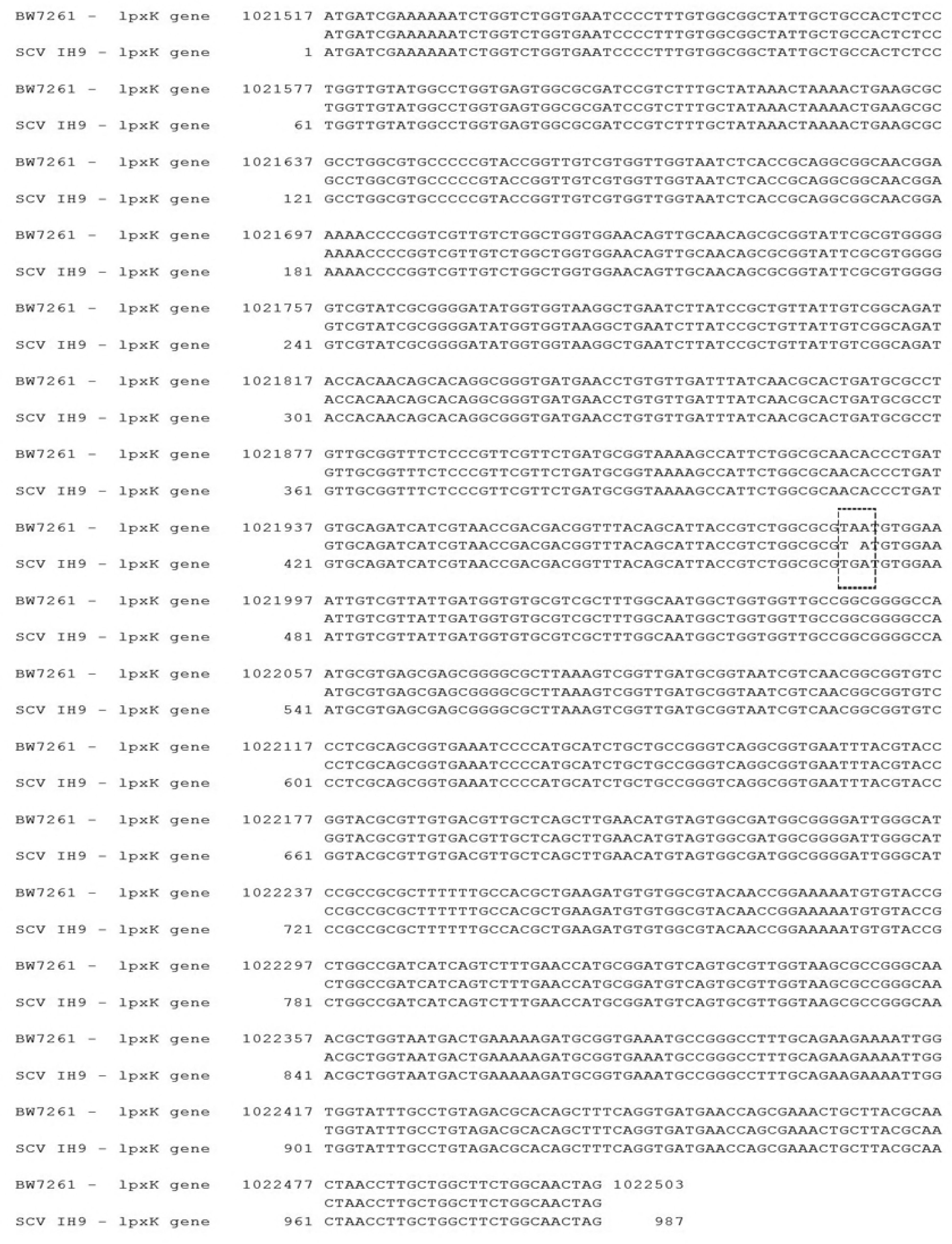
*LpxK* gene nucleotide alignment of SCV IH9 vs. BW7261. SNP is highlighted in dotted box.

**Fig. 13.**
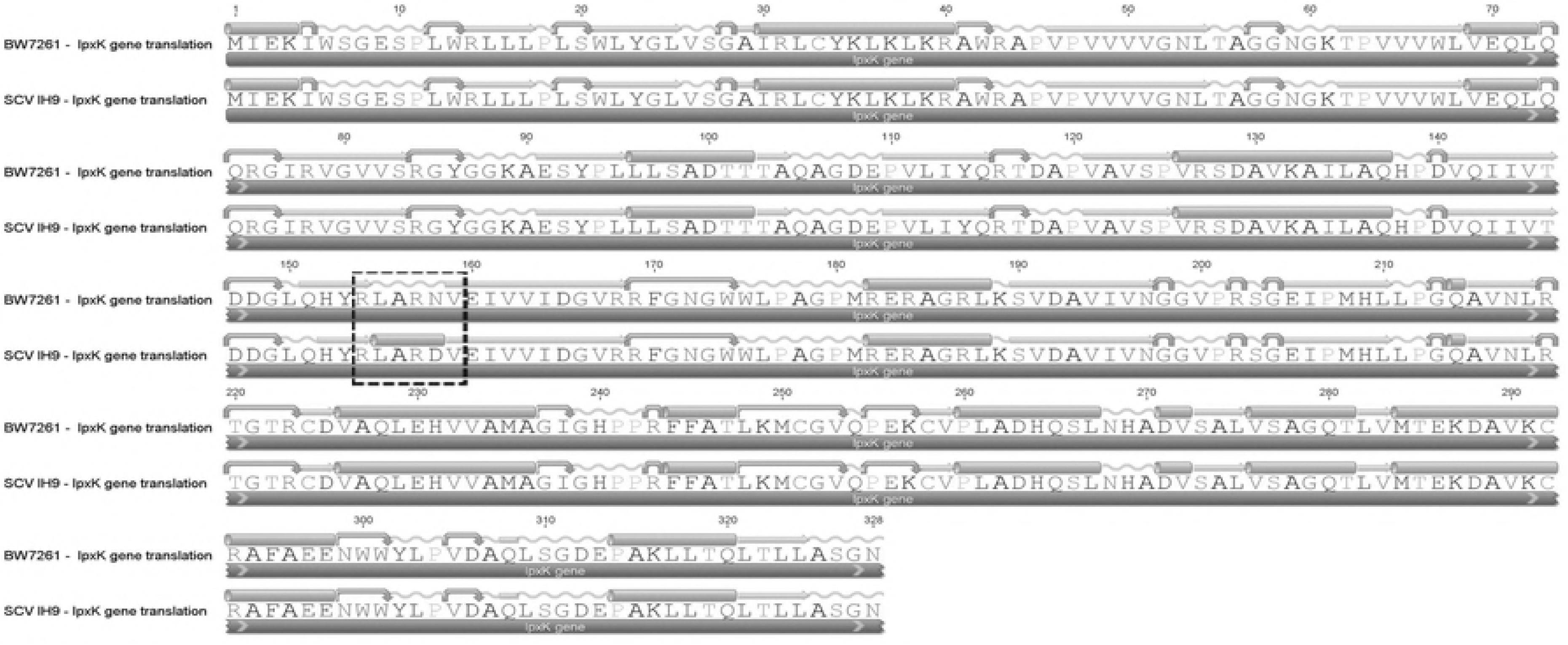
Protein alignment for LpxK in SCV IH9 vs. BW7261. Amino acid substitution is highlighted in dotted box.

## Discussion

SCVs have been identified in scientific literature for at least one hundred years [31]. While the past two decades have seen an expansion in our understanding of SCV physiology, formation, and maintenance, this knowledge concentrates on *Staphylococcus aureus* and recently *Pseudomonas aeruginosa* SCVs [32]. Several articles highlight the association of SCVs with long-term persistent, indolent, chronic and recurring human infections post-surgically [33]. Auxotrophic SCVs have been identified that lack the machinery to synthesize one of three important metabolites; hemin, manadione and thymidine. Recent work identified a SCV of *Escherichia coli* that exhibits auxotrophism for lipoic acid responsible for its small colony size and distinct biochemical features [34].

Our research was prompted by the lack of data profiling the genetic and phenotypic association of *Escherichia coli* SCVs. Although a large body of data exists in published articles, this work mostly offers insight on bacterial physiology and morphological properties of *E. coli* SCVs [35]. Our research revealed over-expression of several genes and gene groups. Specifically, colanic genes involved in biofilm formation and ferric genes (*e.g.* iron transport, iron fixation, etc.) were over-expressed SCV IH9 and may be candidate genes that – on their own – play a major role in SCV formation.

The data presented in this study results from genomic analysis of a SCV IH9 (compared to wild type *E. coli* BW7261). We present strong evidence that (1) several important genes are differentially expressed in SCV IH9 (compared to wild type), (2) nonsense mutations may contribute to the SCV phenotype and (3) SCV IH9 SNPs may influence gene expression of important genes. Importantly, these genetic variants have not been discovered before in *E. coli* SCVs, nor have previous reports detailed the exact pattern of differential expression discovered in this research. Future work will endeavor to elucidate what SNP combination may trigger SCV formation in wild type *E. coli*, perhaps providing a therapeutic target for clinicians to identify future pathogenic SCVs in cultured samples from patients.

## Acknowledgements

We wish to thank Long Pham, Donald Turbiville, Ranjit Singh and Shaveta Anand for their assistance. Finally, we thank St. John’s University for financial support of this research.

